# DisConST: Deciphering Spatial Domains Using Distribution-aware Contrastive Learning for Spatial Transcriptomics

**DOI:** 10.1101/2025.03.13.642300

**Authors:** Peimeng Zhen, Xiaofeng Wang, Han Shu, Jialu Hu, Yongtian Wang, Jiajie Peng, Xuequn Shang, Jing Chen, Tao Wang

## Abstract

Spatial transcriptomics (ST) is a cutting-edge technology that provides comprehensive insights into gene expression patterns from a spatial perspective. A key research focus within this field is spatial domain identification, which is essential for exploring tissue organization, biological development, and disease mechanisms. Although methods have been developed, they still face challenges in modeling the gene expression information together with the spatial locations, resulting in suboptimal accuracy. We introduce DisConST (Distribution-aware Contrastive Learning for Spatial Transcriptomics), a novel deep-learning method designed to improve spatial domain detection within spatial transcriptomics datasets. DisConST addresses key challenges, such as the high dropout rates and the complex integration of spatial and gene expression data, by incorporating contrastive learning strategies that are aware of the underlying data distributions. It employs the zero-inflated negative binomial (ZINB) distribution, along with graph contrastive learning, to generate more informative latent representations. These representations efficiently integrate spatial positions, transcriptomic profiles, and cell-type proportions within spots. We validated DisConST across diverse ST datasets of tissues, organs, and embryos from various sequencing platforms in both normal and disease states. Our results consistently demonstrated that DisConST achieves superior spatial domain recognition accuracy compared to existing state-of-the-art methods. Furthermore, our experiments highlighted the utility of DisConST in advancing research on tissue organization, embryonic development, and tumor immune microenvironment dissection. The source code for DisConST is freely available at https://github.com/Zhenpm/DisConST/.

## Introduction

The spatial organization of cells within tissues is a fundamental aspect of biological systems, influencing everything from developmental processes to disease progression. In tissues such as embryos and tumors, cellular interactions and spatial context play pivotal roles in function and behavior [1]. While conventional single-cell transcriptomics has provided unprecedented insights into cellular gene expression, it falls short of capturing spatial relationships between cells, limiting our understanding of tissue architecture and function. Spatial transcriptomics (ST) is an emerging technology designed to bridge this gap by enabling the comprehensive mapping of gene expression profiles across tissue sections while preserving spatial information [2]. This advancement has opened new avenues for studying cellular organization, tissue structure, and functional heterogeneity in various biological systems. However, a critical task in spatial transcriptomics research is the identification of spatial domains—regions within tissues that exhibit similar gene expression patterns and coherent spatial properties.

Spatial transcriptomics techniques are broadly divided into two categories: next-generation sequencing (NGS)-based approaches and imaging-based methods. NGS-based methods, such as 10X Visium [3], generate spatial gene expression profiles by sequencing transcripts from distinct tissue spots. Imaging-based techniques, such as in situ sequencing (ISS) and in situ hybridization (ISH), offer higher spatial resolution by directly visualizing and quantifying transcripts within tissue sections [4, 5]. Each approach provides distinct advantages in resolution and throughput, but both share the challenge of identifying meaningful spatial domains from high-dimensional and often noisy data.

Traditional clustering algorithms, such as K-means and Louvain, have been applied to categorize spatial domains based solely on gene expression. However, these methods often overlook critical spatial features, leading to discontinuous or inaccurate domain structures. To address these limitations, several advanced techniques have been developed. For example, stLearn [6] normalizes gene expression data using neighborhood information and morphological distance to identify domain structures. SEDR [7] combines deep autoencoders with masked self-supervised learning to create low-dimensional latent representations of gene expression, embedding spatial information with a variational graph autoencoder for domain identification. SpaGCN [8] enhances clustering performance by integrating histological and spatial information, while CCST [9] leverages contrastive learning through randomly constructed graphs to optimize model parameters. BayesSpace [10] employs Bayesian clustering to assign higher weights to physically proximate points, improving model refinement. Similarly, STAGATE [11] utilizes a graph attention encoder to merge spatial and gene expression information, while GraphST [12] enhances representational power through random shuffling of gene expression profiles and contrastive learning. Despite these advancements, existing methods still face challenges due to the high dropout rates in gene expression profiles typical of spatial transcriptomics datasets and difficulties in effectively integrating spatial information, making it challenging to generate highly informative latent representations for accurate domain identification.

To overcome these challenges, we propose DisConST (Distribution-aware Contrastive Learning for Spatial Transcriptomics), a novel method designed to enhance spatial domain detection by leveraging both spatial transcriptomics data and supplementary cell-type proportion information. DisConST employs a deep learning framework, using the zero-inflated negative binomial (ZINB) distribution to model gene expression, which accounts for the sparsity and overdispersion typical of biological data [13, 14]. In addition, DisConST integrates a contrastive learning strategy, optimizing latent representations by aligning gene expression and spatial features through the generation of random graphs and adjacency graphs. These representations are further refined by fusing gene expression data and cell-type proportions through a fully connected encoder, yielding a highly informative latent space for accurate spatial domain detection. We evaluate the performance of DisConST across multiple spatial transcriptomics datasets, spanning both normal and diseased tissues, including organs and embryonic tissues. Our results demonstrate that DisConST consistently outperforms existing state-of-the-art methods in spatial domain identification, providing more accurate and robust insights into tissue organization and function.

## Materials and Methods

This paper presents DisConST, a deep learning method that effectively combines spatial transcriptomics data with cell type proportion data obtained through deconvolution to accurately identify spatial domains. The architecture of DisConST is organized into three main steps, as illustrated in Figure 1. First, DisConST employs an auto-encoder optimized with the zero-inflated negative binomial (ZINB) distribution and graph contrastive learning (GCL) to encode gene expression data and cell type proportion data, respectively. The ZINB distribution is particularly suited for modeling the characteristics of gene expression and cell type proportions, while GCL enhances the discriminative power of the learned representations by leveraging positive and negative pairs. In the second step, DisConST integrates the gene expression and cell type proportion representations obtained from the first step into a unified feature space using a fully connected encoder. Finally, DisConST utilizes a clustering algorithm to predict the spatial domains based on the fused representations.

**Figure 1.**
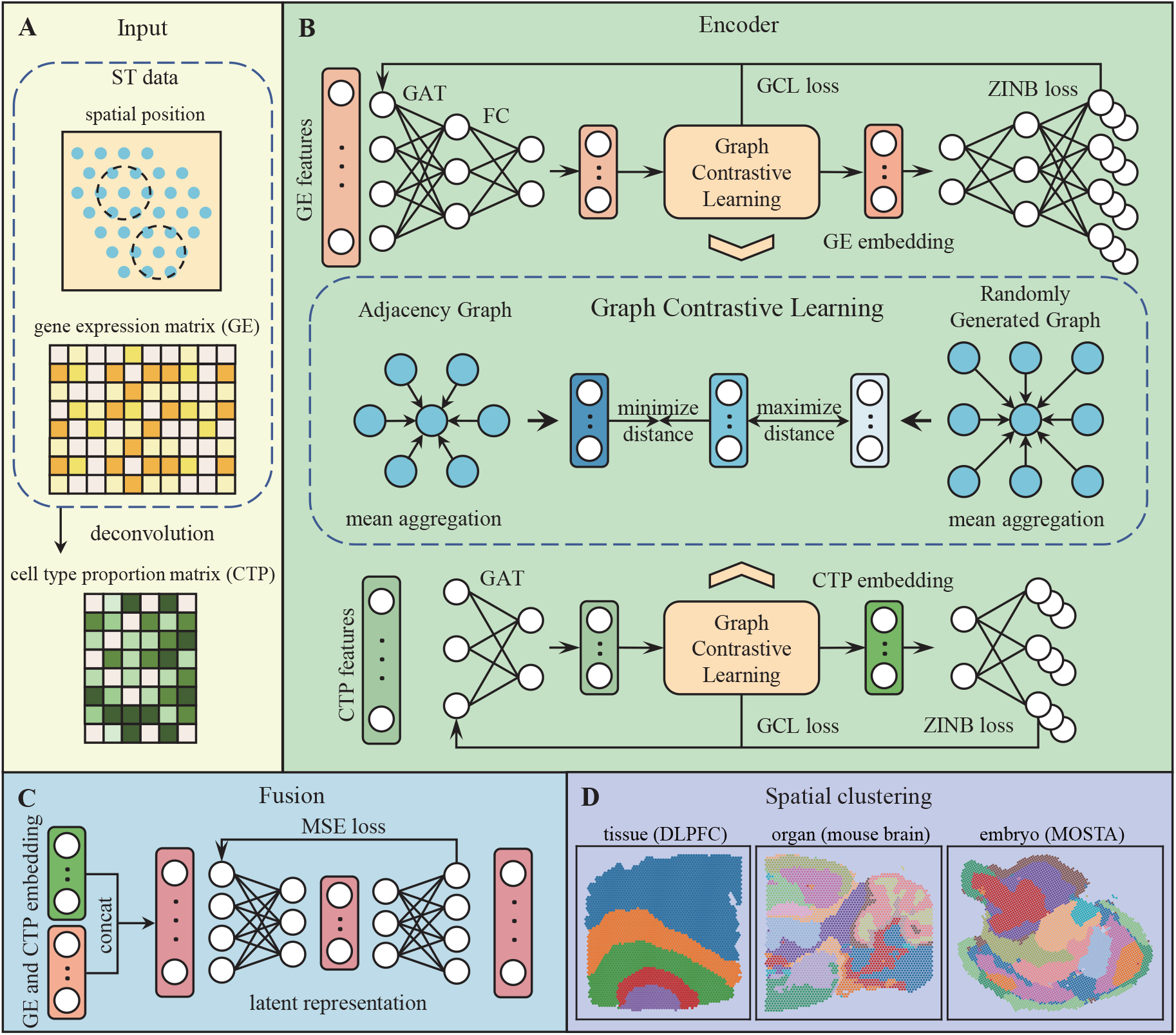
Overview of DisConST workflow. **A**. Input of DisConST. DisConST processes three types of input data: gene expression (GE) data, spatial coordinates, and cell type proportion (CTP) data. It constructs an adjacency graph by incorporating spatial information from the transcriptomics data. The cell type proportions are derived using deconvolution method. **B**. Encoder architecture of DisConST. DisConST uses a dual-encoder architecture to encode both the GE data and CTP data with a similar structure. The encoder is optimized with a combination of Zero-Inflated Negative Binomial (ZINB) loss and graph contrastive learning loss. **C**. Representation fusion. DisConST integrates the GE and CTP embeddings through a fully connected encoder to generate the final latent representation by concatenating these features. **D**. Spatial clustering for domain identification. DisConST applies spatial clustering techniques to recognize spatial domains in tissues, organs, and embryos.

### Data preprocessing

For the dataset used in this study, we first selected a set of n highly variable genes (HVGs), typically set to 3000 for datasets with a sufficient number of sequenced genes. This selection was performed using the scanpy package [15]. The selected genes underwent normalization and log transformation, which provided the initial features of gene expression for DisConST, referred to as GE features. Additionally, we processed the spatial transcriptomics data using deconvolution tools such as SPACEL [16] and Tangram [17] to obtain cell type proportions for each spot. To be noted, if the spatial transcriptomics data is at single-cell resolution, the cell type information will not be considered. The log-transformed cell type proportions were then utilized as the second set of initial features in DisConST, termed CTP features.

### Spatial adjacency graph construction

The spatial transcriptomics data provides the coordinates for each spot, which serve as the basis for constructing a spatial adjacency graph (SAG). In the SAG, each spot is represented as a node, with edges connecting nodes that are in close proximity. Specifically, we select the k nearest nodes for each node to construct the SAG (default k=5). We denote the adjacency matrix as *A*, where if there is an edge between nodes *i* and *j*, then *A*_*ij*_ is set to 1; otherwise, *A*_*ij*_ is set to 0.

### Graph encoder

To obtain the latent representation of spots, DisConST utilizes the spatial adjacency graph (SAG), along with the preprocessed gene expression matrix and cell type proportion matrix, as inputs to the graph encoder. For encoding the gene expression data, we employ a two-layer neural network, where the first layer is a graph attention (GAT) layer [18], followed by a fully connected (FC) layer. The graph attention layer incorporates an attention mechanism that adaptively assigns weights to a node’s neighbors, enhancing feature aggregation by emphasizing contributions from similar neighbors while diminishing those from dissimilar ones. The computation of attention weights occurs in two steps, beginning with the calculation of attention scores for the node pair (*i, j*) as follows:

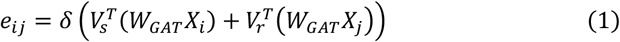

where *X*_*i*_ and *X*_*j*_ are the initial representation of node *i, j*, specifically, the input gene expression data. *V*_*r*_ and *V*_*r* are trainable parameters, while *W*_*GAT*_ serves as the trainable matrix for feature extraction in GAT. δ is the sigomid activation function. The second step is to normalize the attention score to obtain the attention coefficient, which can be described as follow:

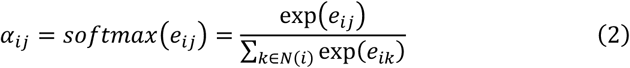

where α_*ij*_ is the attention coefficient of node pair (*i, j*). *N*(*i*) is the set of all neighbors of node *i*. Based on the attention coefficient, GAT aggregates the representation of the central node from neighboring nodes:

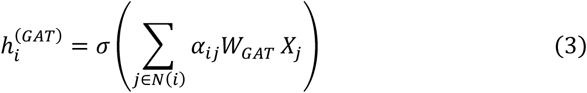

where 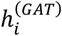 represents the first layer representation of spot *i* obtained from GAT layer, and σ is the ELU activation function. To further extract latent representations of gene expression features for each node, DisConST employs a fully connected layer to reduce the output dimension of the GAT layer:

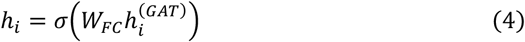

Here, *h*_*i*_ represents the latent representation of spot *i* obtained from the fully connected layer, σ denotes the ELU activation function, and *W*_*FC*_ is a trainable parameter.

For the cell type proportion data, the encoding process is similar, but due to the relatively low number of cell types in many datasets, we use a single-layer graph attention network (GAT) for encoding without a subsequent fully connected layer.

### Optimization Strategy

The loss function of DisConST is divided into two parts: ZINB loss and graph contrastive learning loss.

#### Zero-Inflated negative binomial distribution-based loss

The ZINB distribution is commonly used to describe the characteristics of single-cell data [14]. Our experiments have shown that the ZINB distribution performs well in spatial transcriptomics data. This distribution combines two probability distributions to better fit data with a large number of zeros. Its probability mass function can be expressed as follows:

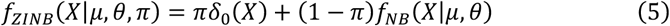

where *π* is the dropout probability (the proportion of zeros). δ_0_ is a one-point distribution that takes the value 1 at 0 and 0 at all other positions. fN B is the probability mass function of the negative binomial (NB) distribution, expressed as:

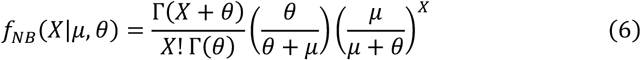

where *μ* and *θ* represent the mean and dispersion parameters of the NB distribution, respectively. To construct the ZINB loss, we design a special decoder to obtain the three parameters of the ZINB distribution. The decoder and encoder are symmetrical structures. For a two-layer encoder, we construct a two-layer decoder, with the first layer defined as follows:

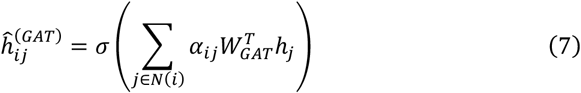

Corresponding to the GAT layer, the first layer decoder uses the same parameters, where 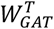 is the transpose of *W*_*GAT*_ and 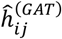 is the GAT decoding representation of node *i*. To obtain the three parameters of the ZINB distribution, the second layer has three parts, defined as follows:

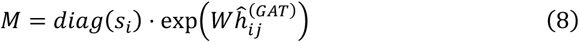

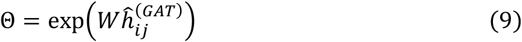

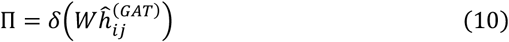

In the second layer, *M*, Θ, and Π represent matrix estimates of mean, dispersion, and dropout probability, respectively. For the mean and dispersion, we choose the exponential activation function, as both parameters are non-negative values. For the additional coefficient Π, the activation function δ is sigmoid, representing the dropout probability. As the dropout probability ranges from 0 to 1, sigmoid is an appropriate choice. *W* is a shared trainable parameter, and *s*_*i*_ is the size factor of spot *i*, defined as:

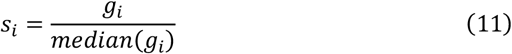

where *g*_*i*_ is the total gene expression of spot *i*, representing the relative gene expression of each spot. Similar to the encoder, for datasets with relatively few cell types, we also have only one decoder layer, specifically defined as follows:

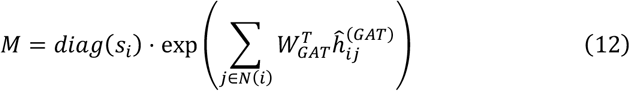

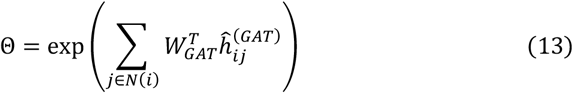

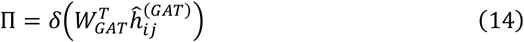

Finally, we use the negative logarithmic likelihood function to calculate the ZINB loss. Specifically, the ZINB loss is defined as follows:

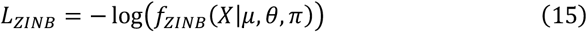

#### Graph contrastive learning-based loss

Although the graph auto-encoder can integrate features from neighboring nodes to learn latent representations, it may lead to over-smoothed representations that obscure distinct biological features, complicating effective clustering. To enhance the discriminative power of the latent representation, we employ a graph contrastive learning process.

Firstly, for each node *i*, we randomly select *l* nodes (excluding node *i*). Due to the significant gap between the total number of nodes in the graph and the number of neighbors of node *i*, these nodes are usually not neighbors of node *i* but rather located far from it. This gives us a randomly generated graph (RGG) *G*′ = (*V*′, *E*′). We then aggregate the neighbors of each node in the SAG *G* and RRG *G*′ through mean aggregation, defined as follows:

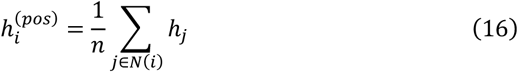

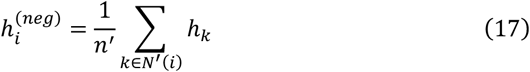

We denote *h*_*i*_ as the representation of anchor spot *i*, where 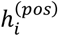 is the representation obtained by aggregating the neighbors of node *i* from the SAG, forming a positive pair with *h*, while 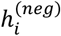 is the representation obtained from the RRG, forming a negative pair with *h*_*i*_. *N*(*i*) and *N*′(*i*) are the sets of all neighbors of node *i* in SAG and RRG, respectively, and *n* and *n*′ are the numbers of neighbors corresponding to node *i* in the two graphs.

To achieve contrastive learning by minimizing the Euclidean distance between positive pairs and maximizing the distance between negative pairs, the loss function is specifically formulated as follows:

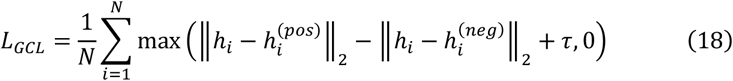

where τ is used to enforce a margin between the distances of positive and negative pairs, with a default value of 1. *N* represents the total number of spots, as each spot corresponds to one positive pair and one negative pair.

The final loss is represented as:

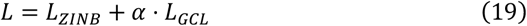

where α is the coefficient for the GCL loss, with a default value of 0.5.

### Feature fusion

To jointly utilize the latent representations obtained from gene expression data and cell type proportion data for subsequent tasks, we employ a fully connected encoder to fuse the two representations as follows:

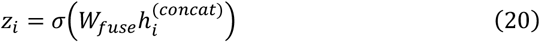

where *z*_*i*_ represents the final latent representation of spot *i*, and 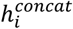 is the representation obtained by concatenating the latent representations of gene expression and cell type proportions. Subsequently, DisConST utilizes a fully connected decoder to decode the fused representation. The fusion encoder employs mean squared error loss to describe the reconstruction effect:

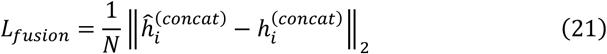

where 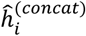 represents the decoded reconstruction of 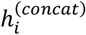.

### Clustering and refinement

We utilize the Gaussian mixture model from the Mclust R package [19] version 6.0.0 to obtain clustering results for the latent representations learned by DisConST. The number of clusters is based on the ground truth labels. To enhance clustering accuracy, we implement a refinement step. For each spot *i*, we define a radius *r* (default value of 50) within which all spots are considered neighbors of spot *i*. We then reassign spot *i* to the domain with the most common label among its neighbors. Notably, this refinement process may not be suitable for fine-grained datasets, such as those from the mouse olfactory bulb, brain, and embryos. In this study, we applied this refinement step only to the human brain DLPFC and the human breast cancer datasets.

### Benchmark

We compared DisConST with seven state-of-the-art methods, including stLearn [6], SEDR [7], SpaGCN [8], CCST [9], BayesSpace [10], STAGATE [11], and GraphST [12]. We used the Adjusted Rand Index (ARI) [20] to evaluate the clustering results, defined as follows:

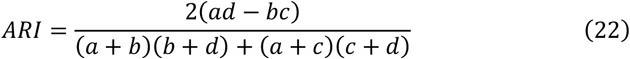

In this equation, *a* denotes the number of spot pairs that are in the same cluster under both actual and experimental conditions. *b* represents the number of spot pairs that are in the same cluster in real situations but not in the same cluster in experimental situations. *c* indicates the number of spot pairs that are not in the same cluster in real situations but are in the same cluster in experimental situations. Finally, *d* stands for the number of spot pairs that are not in the same cluster in either real or experimental situations. The ARI ranges from [-1, 1], with higher values signifying greater congruence between the clustering result and the ground truth labels, thereby indicating better clustering performance.

## Data Availability

In our work, we employed six distinct datasets derived from various sequencing platforms, including 10X Visium, Stereo-seq, seqFISH, and Spatial Transcriptomics. The first dataset consisted of twelve DLPFC tissue sections obtained from 10X Visium, featuring between 3,460 and 4,789 spots, encompassing 33,538 genes. These data, along with manual annotations, are available at http://research.libd.org/spatialLIBD/. The second set comprised mouse olfactory bulb slices acquired with Stereo-seq and Spatial Transcriptomics, with spot counts of 10,000 and 282, respectively. These ST data are available at https://github.com/hannshu/st_datasets. The third cohort included mouse embryo slices derived from Stereo-seq and seqFISH (https://crukci.shinyapps.io/SpatialMouseAtlas/), containing 5913 and 19416 spots each. Additionally, four mouse brain tissue sections from the 10X Visium platform (https://www.10xgenomics.com/resources/datasets) were utilized, exhibiting spot numbers ranging from 2,696 to 3,353. The fifth dataset featured human breast cancer tissue samples from 10X Visium (https://www.10xgenomics.com/datasets/human-breast-cancer-block-a-section-1-1-standard-1-1-0), with a spot count of 3,798. Lastly, we employed four Mouse Organogenesis Spatiotemporal Transcriptomic Atlas datasets (E9.5-E12.5) from the Stereo-seq platform (https://db.cngb.org/stomics/mosta/download/), with spot counts ranging from 5,059 to 27,455.

For datasets from 10X Visium, where the resolution is multi-cell per spot, we used cell type proportions as supplementary data, necessitating single-cell transcriptomics data for deconvolution. First, single-nucleus transcriptomics data across multiple human cortical areas are available at https://portal.brain-map.org/atlases-and-data/rnaseq/human-multiple-cortical-areas-smart-seq. Second, single-cell transcriptomics data for human breast cancer is available at https://singlecell.broadinstitute.org/single_cell/study/SCP1039. Finally, single-cell transcriptomics data of the mouse whole brain is available at mousebrain.org/adolescent/downloads.html.

## Results

### Overview of DisConST workflow

In this work, we propose a graph neural network-based method, DisConST, for representation learning and spatial domain identification of spatial transcriptomics data (Figure 1). DisConST encodes the spot-level representations by considering both the gene expressions and the cell types within a spot, simultaneously. Specifically, DisConST first constructs a Spatial Adjacency Graph (SAG) using the spatial locations of spots, and then utilizes a graph neural network to encode gene expression profiles and cell type proportions separately. The encoder consists of Graph Attention Network (GAT) and Fully Connected (FC) layers for feature extraction, which adaptively integrates features between the central spot and its spatially nearby spots. During the training process, the model is optimized by two strategies. First, the Zero-Inflated Negative Binomial (ZINB) distribution is used in modeling the input features to address excess zeros and overdispersion simultaneously. Second, DisConST uses graph contrastive learning to enhance the discrimination of the latent representations (See Materials and Methods). After separately encoding gene expression and cell type information, DisConST uses an additional fully connected layer to fuse the two representations. The final integrated representation is then used for spatial domain identification based on unsupervised clustering.

### DisConST enables accurate spatial domain identification on human dorsolateral prefrontal cortex

In order to evaluate the performance of DisConST, we used 12 slices of the human dorsolateral prefrontal cortex (DLPFC) from 10X Visium sequencing technology for spatial domain identification [21]. The 12 slices of DLPFC have been manually annotated as 6 different layers and white matter (WM), which serve as the ground truth for evaluation. We compared DisConST with 7 state-of-the-art methods, namely stlearn [6], SEDR [7], SpaGCN [8], CCST [9], BayesSpace [10], STAGATE [11], GraphST [12]. And the performance was evaluated using the Adjusted Rand Index (ARI) metric [20].

Figure 2A compares the performance of DisConST with seven state-of-the-art methods across 12 DLPFC slices. DisConST achieves an average Adjusted Rand Index (ARI) of 0.62, with a peak ARI of 0.86. Among the competitors, STAGATE and GraphST perform best, with average ARIs of 0.50 and 0.51, respectively. Additionally, we use the first slice (#151507) to further illustrate the clustering accuracy of DisConST compared to other methods (Figure 2B, C). Notably, only DisConST accurately captures the structure of layers 2 and 3, which are positioned on both sides of layer 1. Moreover, stLearn, SEDR, and SpaGCN fail to recognize the hierarchical structure from layers 4 to 6, resulting in relatively disorganized domain partitions. CCST incorrectly merges these three distinct layers into a single domain.

**Figure 2.**
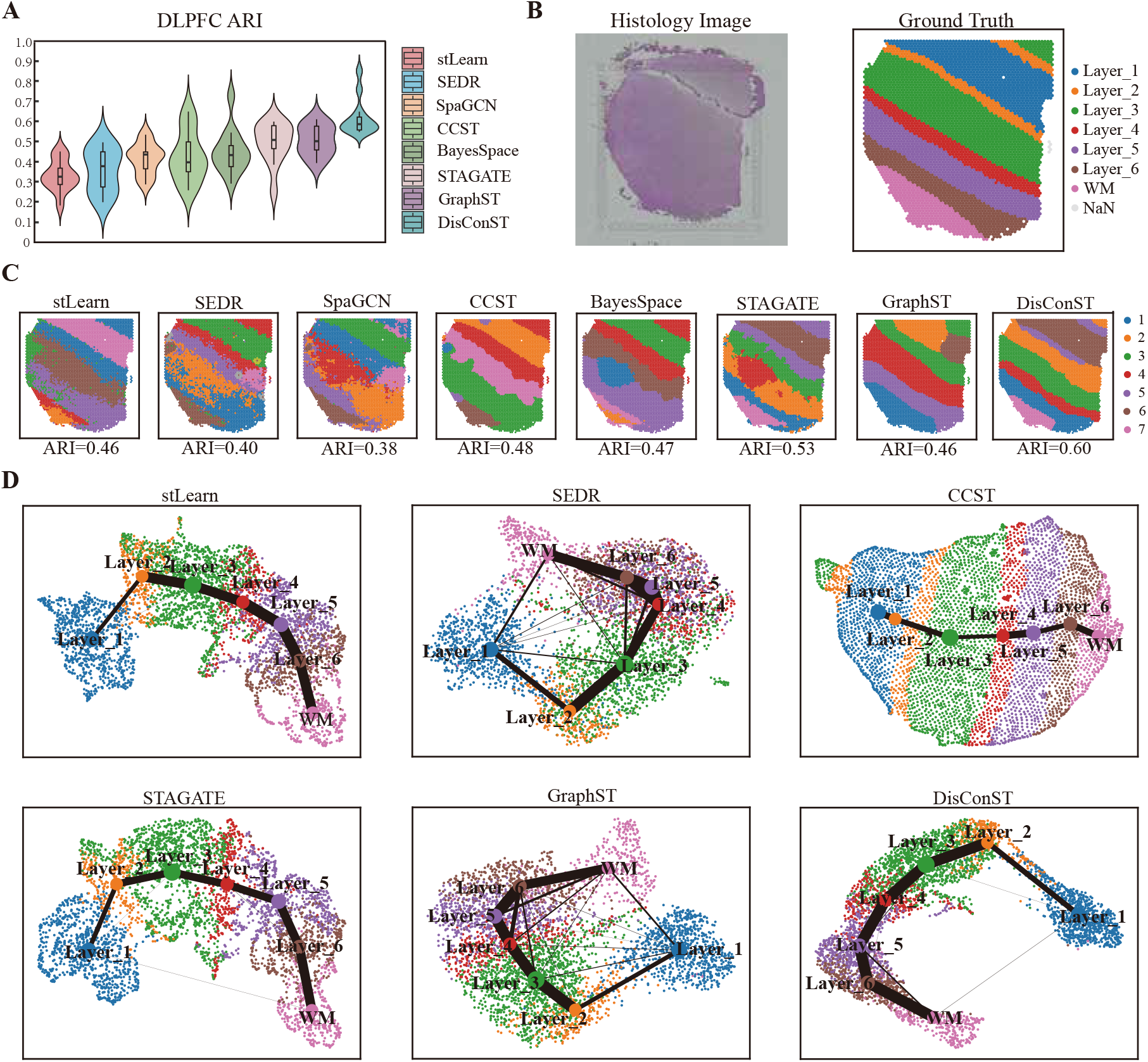
DisConST significantly improves the accuracy of spatial domain recognition on human dorsolateral prefrontal cortex (DLPFC) spatial transcriptomics dataset. **A**. the violin plot shows the Adjusted Rand Index (ARI) to summarize the accuracy of each method in all 12 slices of the DLPFC dataset. The box plot is embedded in the violin plot. The boxplot’s center line, box limits, and whiskers denote the median, upper and lower quartiles, and 1.5× interquartile range, respectively. **B**. The histology image and ground truth with six cortex layers and white matter (WM) of DLPFC slice #151507. **C**. Visualization of spatial domain detection of eight methods, stLearn, SEDR, SpaGCN, CCST, BayesSpace, STAGATE, GraphST, and DisConST on DLPFC slice #151507. **D**. UMAP plots and corresponding PAGA spots of latent representations obtained from six methods, stLearn, SEDR, CCST, STAGATE, GraphST, and DisConST. Notably, as end-to-end methods, SpaGCN and BayesSpace do not provide latent representations, causing inability to visualize UMAP and PAGA plots.

While BayesSpace and STAGATE successfully identify the hierarchical arrangement of layers, they incorrectly produce an elliptical domain in this region. Lastly, GraphST erroneously splits layer 1 into two separate domains. In summary, DisConST is the only method that correctly identifies the overall layer structure. A comprehensive performance summary for all 12 slices can be found in Supplementary Figure S1 and Supplementary Table 1.

Additionally, we generated UMAP visualization plots for DisConST and five other methods using DLPFC slice #151507 to further evaluate the expressiveness of their latent representations. The underlying assumption is that effective latent representations should accurately preserve the original relative positions of spatial domains. As shown in Figure 2D, the UMAP visualizations for SEDR and GraphST fail to capture the hierarchical structure among the six cortical layers and the white matter, while other methods demonstrate clearer layer distribution patterns. Notably, compared to the other five methods, DisConST not only distinguishes each domain more effectively but also exhibits tighter clustering of spots within the same domain. To further validate the strength of our latent representations, we applied the PAGA algorithm [22] for trajectory inference. The trajectory plots are overlaid on the UMAP plots in Figure 2D. DisConST shows a nearly linear developmental trajectory from the first layer to the white matter, with higher similarity between adjacent layers. In contrast, the trajectories inferred from SEDR and GraphST reveal little to no discernible linear relationships.

### DisConST demonstrates robust performance across different sequencing platforms

As spatial transcriptomics technology evolves, various sequencing techniques with different resolution levels have emerged. To assess the performance of DisConST across different platforms, we conducted spatial domain identification using multiple sequencing technologies, including Stereo-seq, Spatial Transcriptomics, and seqFISH. For this experiment, we used spatial transcriptomics (ST) data from mouse olfactory bulb tissue sequenced by Stereo-seq [23] and Spatial Transcriptomics [3]. Additionally, we analyzed developmental mouse embryos using data from Stereo-seq [24] and seqFISH [25]. Both Stereo-seq and seqFISH offer single-cell resolution, whereas Spatial Transcriptomics has a lower resolution, with each spot covering a diameter of approximately 100 μm.

As shown in Figure 3A, DisConST significantly outperforms the two leading comparison models, STAGATE and GraphST, in identifying spatial domains within mouse olfactory bulb tissue using both the Stereo-seq and Spatial Transcriptomics platforms. DisConST achieves an ARI above 0.5 for data from both platforms, while GraphST struggles to identify domain structures, resulting in mixed domain distributions. Additionally, as shown in Figure S2, DisConST more effectively reveals the hierarchical structure of the mouse olfactory bulb compared to STAGATE and GraphST. To further validate DisConST’s cross-platform capability, we extended our experiments to developing mouse embryos, applying three clustering methods to seqFISH data from stages E8.5 to E8.75, and to Stereo-seq data from stage E9.5. Figure 3B demonstrates that, despite variations in developmental stages and tissue sequencing techniques, DisConST consistently outperforms the other two methods. In summary, DisConST maintains robust spatial domain detection performance across different resolutions and sequencing platforms.

**Figure 3.**
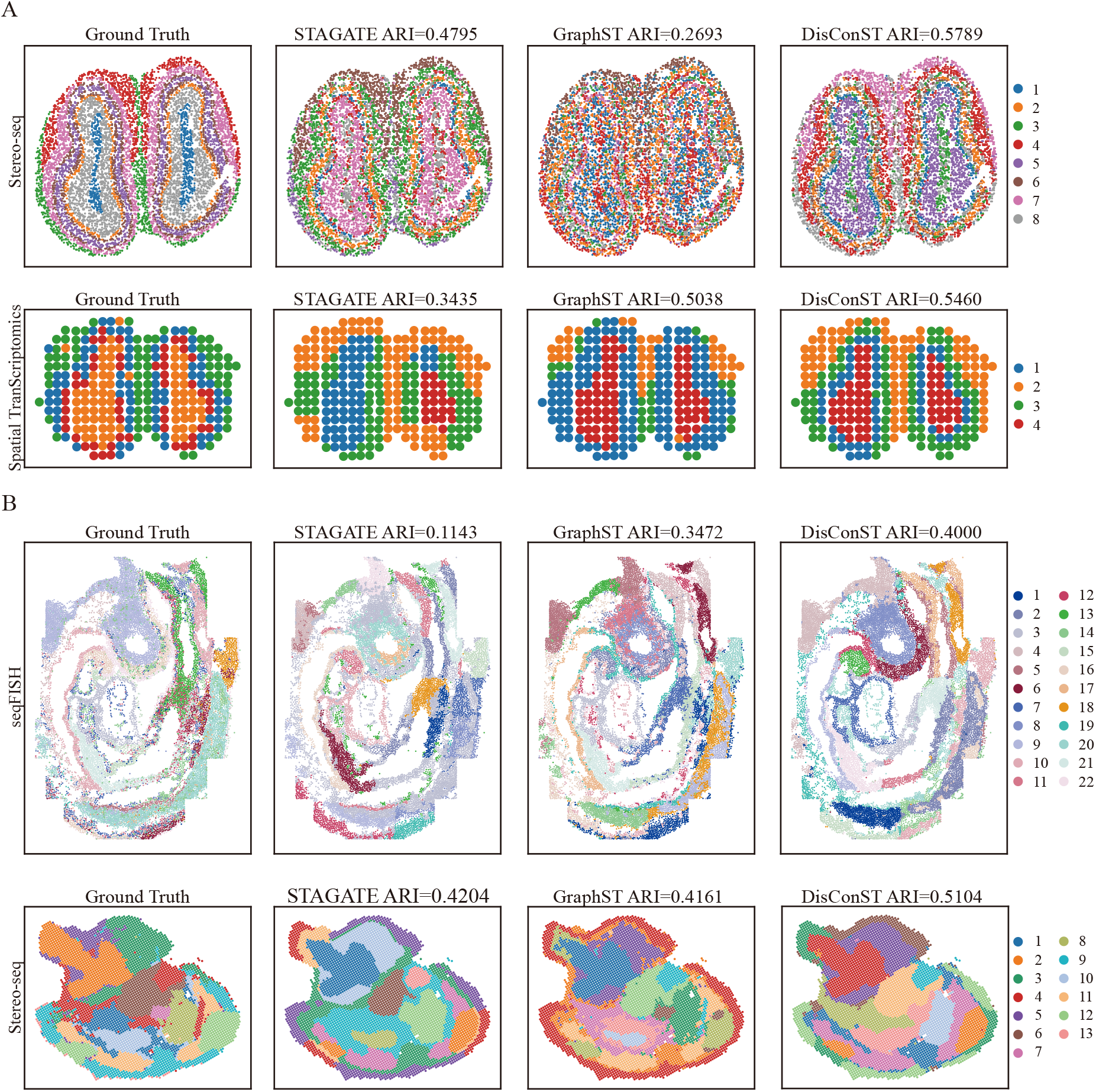
DisConST demonstrates robust performance across different sequencing platforms. **A**. Ground truth and spatial domains identified by STAGATE, GraphST, and DisConST for mouse olfactory bulb tissue, sequenced using Stereo-seq and Spatial Transcriptomics. **B**. Ground truth and spatial domains identified by STAGATE, GraphST, and DisConST for the mouse organogenesis spatiotemporal transcriptomic atlas, sequenced using seqFISH and Stereo-seq. The ARI scores are indicated for each plot.

### DisConST effectively identifies the organizational structure of the mouse brain

To demonstrate DisConST’s ability to identify complex organizational structures, we analyzed two sections of mouse brain slices using 10X Visium technology. As shown in Figure 4A and B, each section is divided into anterior and posterior sagittal portions, forming the complete brain structure. In Figure 4C, we visualized the ground truth domain distribution from STomicsDB [26], alongside spatial domain identification results from three methods applied to the two mouse brain slices. DisConST consistently outperformed STAGATE and GraphST in terms of ARI scores across both slices. Notably, DisConST accurately identified the small olfactory bulb and the complex cerebellar cortex tissues (highlighted with green and blue boxes, respectively). Moreover, the domains corresponding to the midbrain (MB), medulla (MY), and hippocampus (HPF) regions, marked as MB, MY1/2, and HPF1/2 in Figure 4C, align well with the ground truth.

**Figure 4.**
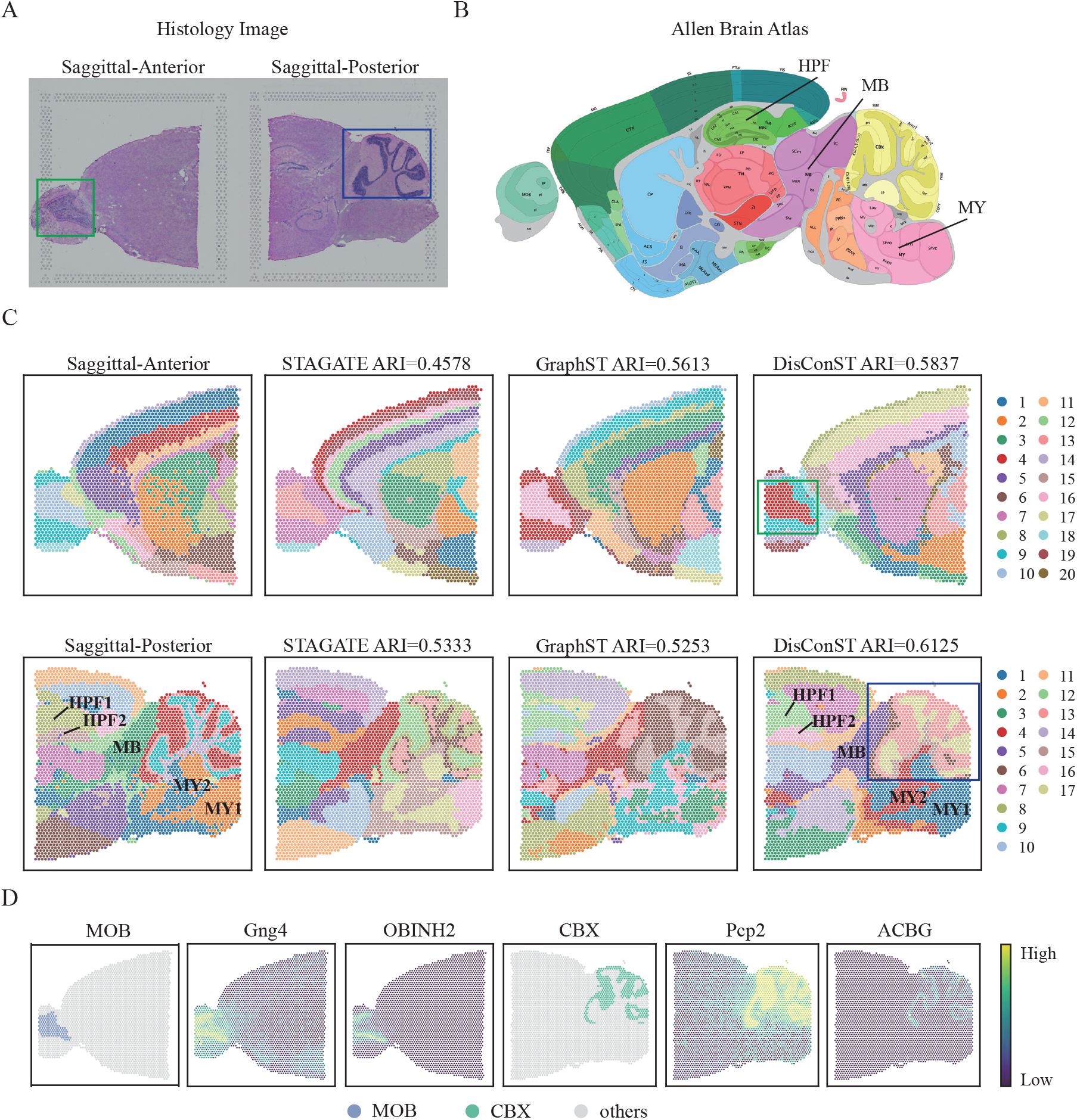
DisConST effectively identifies tissue structure in the mouse brain. **A**. Histological images of sagittal-anterior and sagittal-posterior sections of the mouse brain. **B**. The Allen Brain Atlas for sagittal mouse brain at position 121. **C**. Visualization of spatial domain identification by STAGATE, GraphST, and DisConST in the sagittal-anterior and sagittal-posterior sections, alongside corresponding ground truth. **D**. Marker genes and highly prevalent cell types in the mouse olfactory bulb (MOB) and cerebellar cortex (CBX).

Furthermore, the spatial expression patterns of marker genes and the distribution of corresponding cell types further validate the accuracy of the tissue identification. As shown in Figure 4D, the protein-coding gene *Gng4* exhibits high expression in the mouse olfactory bulb tissue [27]. Similarly, OBINH2, a kind of olfactory inhibitory neurons, is also significantly concentrated in the same region [28]. These observations align well with the olfactory bulb region identified by DisConST. Additionally, *Pcp2*, a gene encoding Purkinje cell protein 2 [29], is highly expressed in Purkinje cells of the mammalian cerebellum, while ACBG, cerebellar Bergmann glial cells [28], is distinctly found in the cerebellum. This supports DisConST’s accurate identification of the cerebellar region. The alignment of marker gene expression and cell type distribution with DisConST’s results further confirms the precision of its tissue structure identification. The outcomes for the second section, which reflect similar findings, are detailed in Supplementary Figure S3.

### DisConST accurately identifies the tissue structures of mouse embryo

Research on embryonic development is a critical application of spatial transcriptomics technology. By analyzing the spatial distribution of transcripts within embryos, we can gain valuable insights into tissue and organ formation and differentiation, providing a deeper understanding of these processes from a spatial perspective. Specifically, spatial transcriptomics at various developmental stages enables us to observe the dynamic progression of embryonic development. To evaluate DisConST’s effectiveness in identifying organ and tissue structures in developing embryos, we applied it to the Mouse Organogenesis Spatiotemporal Transcriptomic Atlas (MOSTA) data from four developmental stages (E9.5, E10.5, E11.5, and E12.5), generated using Stereo-seq sequencing technology [24]. As illustrated in Figure 5A, manual annotations capture the formation and developmental processes of multiple organs. As shown in Figure 5B, DisConST accurately identifies the overall structures of embryonic tissues and organs across all four slices, achieving an average ARI of 0.4, which outperforms STAGATE and GraphST, whose ARI scores are 0.356 and 0.325, respectively (Figure 5C).

**Figure 5.**
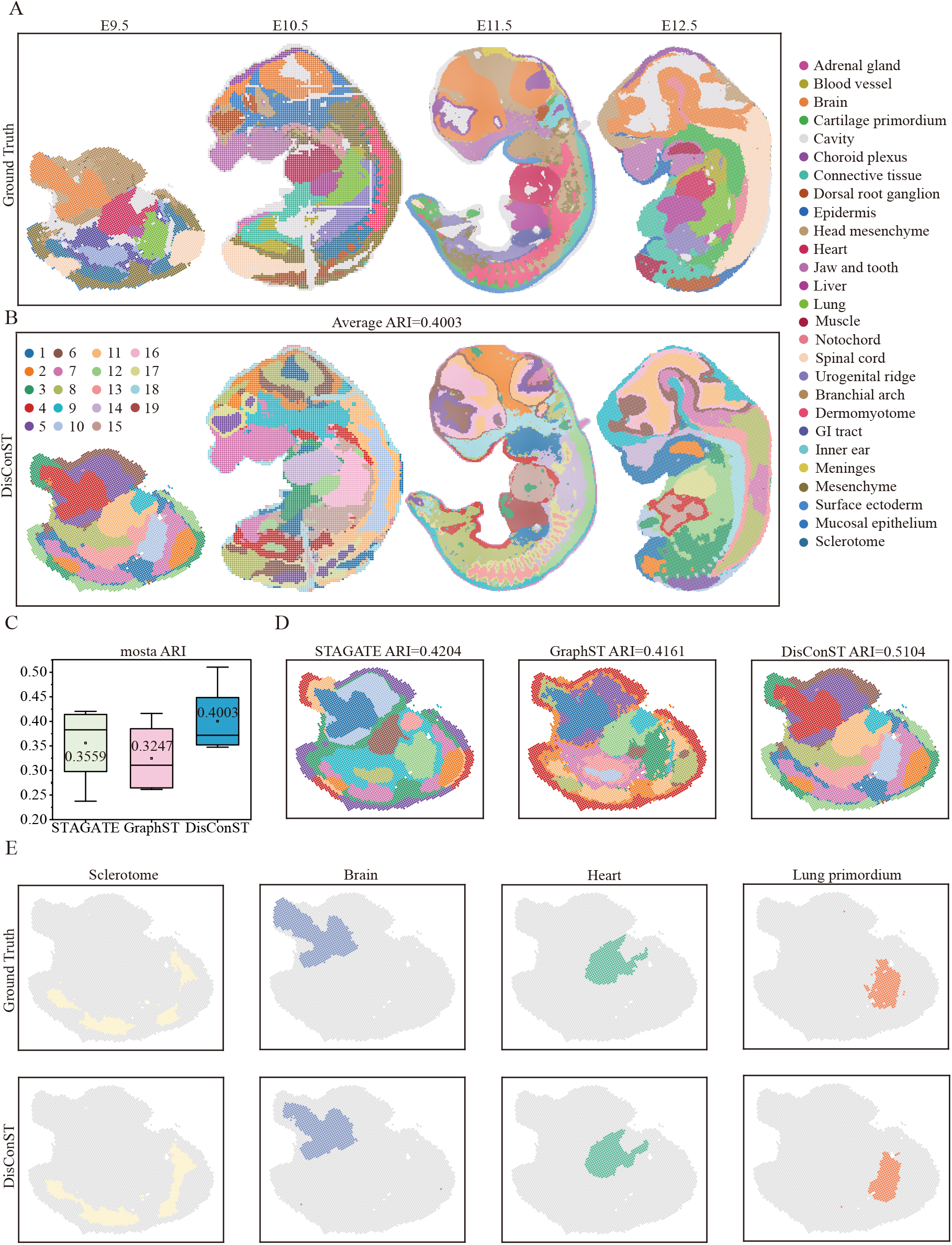
DisConST accurately identifies tissue structures in mouse embryos. **A**. Ground truth of the Mouse Organogenesis Spatiotemporal Transcriptomic Atlas (MOSTA) at four different developmental stages (E9.5–E12.5). **B**. Spatial domains detected by DisConST across the four MOSTA datasets. **C**. Box plots of ARI scores for spatial domains identified by STAGATE, GraphST, and DisConST on the MOSTA datasets. The center line, box limits, and whiskers represent the median, upper and lower quartiles, and 1.5× interquartile range, respectively. The average ARI score is also indicated within the box plot. **D**. Visualization of spatial domains detected by the three methods on the MOSTA E9.5 dataset. **E**. Spatial domains detected by DisConST with corresponding annotations for four tissues or organs: sclerotome, brain, heart, and lung primordium.

Using stage E9.5 as an example, Figure 5D highlights DisConST’s superior performance, achieving an ARI score of 0.51, approximately 25% higher than STAGATE and GraphST. DisConST successfully identifies domains with well-defined boundaries and organ shapes that closely align with the ground truth, including the sclerotome, brain, heart, and lung primordium, as shown in Figure 5E. In contrast, GraphST produces blurred domain boundaries, and STAGATE inaccurately captures the shapes of key organs, such as the heart. For replication purposes, the performance was also evaluated in the other four slices at stage E9.5, as shown in Supplementary Figure S4.

The clustering results for STAGATE and GraphST at other stages are presented in Supplementary Figure S5. A comparative analysis of spatial domain identification across the four stages of embryonic development further demonstrates the superior performance of DisConST in cross-stage identification of organizational structures. As illustrated in Supplementary Figure S6, DisConST consistently identified the heart structure in all four stages. Additionally, despite changes in the jaw and tooth structures observed in the last three stages, DisConST still achieved accurate identification. Moreover, DisConST detected the appearance and structure of muscle tissue, which emerged at the E12.5 stage. In summary, DisConST effectively reveals the early embryonic process of organ formation and tissue structuring, offering valuable insights for embryonic development research.

### DisConST effectively dissects the tumor immune microenvironment

Spatial transcriptomics plays a crucial role in tumor research by preserving the spatial distribution of tumor domains, thereby providing valuable insights into the tumor microenvironment. To evaluate the effectiveness of DisConST in analyzing the immune microenvironment, we applied it to spatial transcriptomics data from breast cancer using 10X Visium technology. This dataset exhibited high heterogeneity and complex domain structures [30], contrasting with the distinct hierarchical organization and high similarity between layers observed in the DLPFC dataset.

As illustrated in Figures 6A and B, the histological image is divided into 20 domains based on pathological features. To highlight the accuracy of DisConST, we compared its spatial domain identification results with those of STAGATE and GraphST. Figure 6C shows that after refining the identification results from the three methods, DisConST achieved an ARI approximately 25% higher than the other two methods. Additionally, the identification of structures within multiple diseased areas, such as DCIS/LCIS and IDC, strongly aligns with the ground truth. Notably, DisConST accurately identified specific cancerous regions, including DCIS/LCIS 1 (ductal carcinoma in situ/lobular carcinoma in situ) and IDC 4 (infiltrating ductal carcinoma), marked with green and yellow boxes, respectively. In contrast, both GraphST and STAGATE failed to accurately identify the structure of the DCIS/LCIS 1 area, missing the outer tumor edge in region 3, and incorrectly segmented IDC 4 into two separate domains. These results demonstrate that DisConST can accurately delineate the distribution of tumor domains.

**Figure 6.**
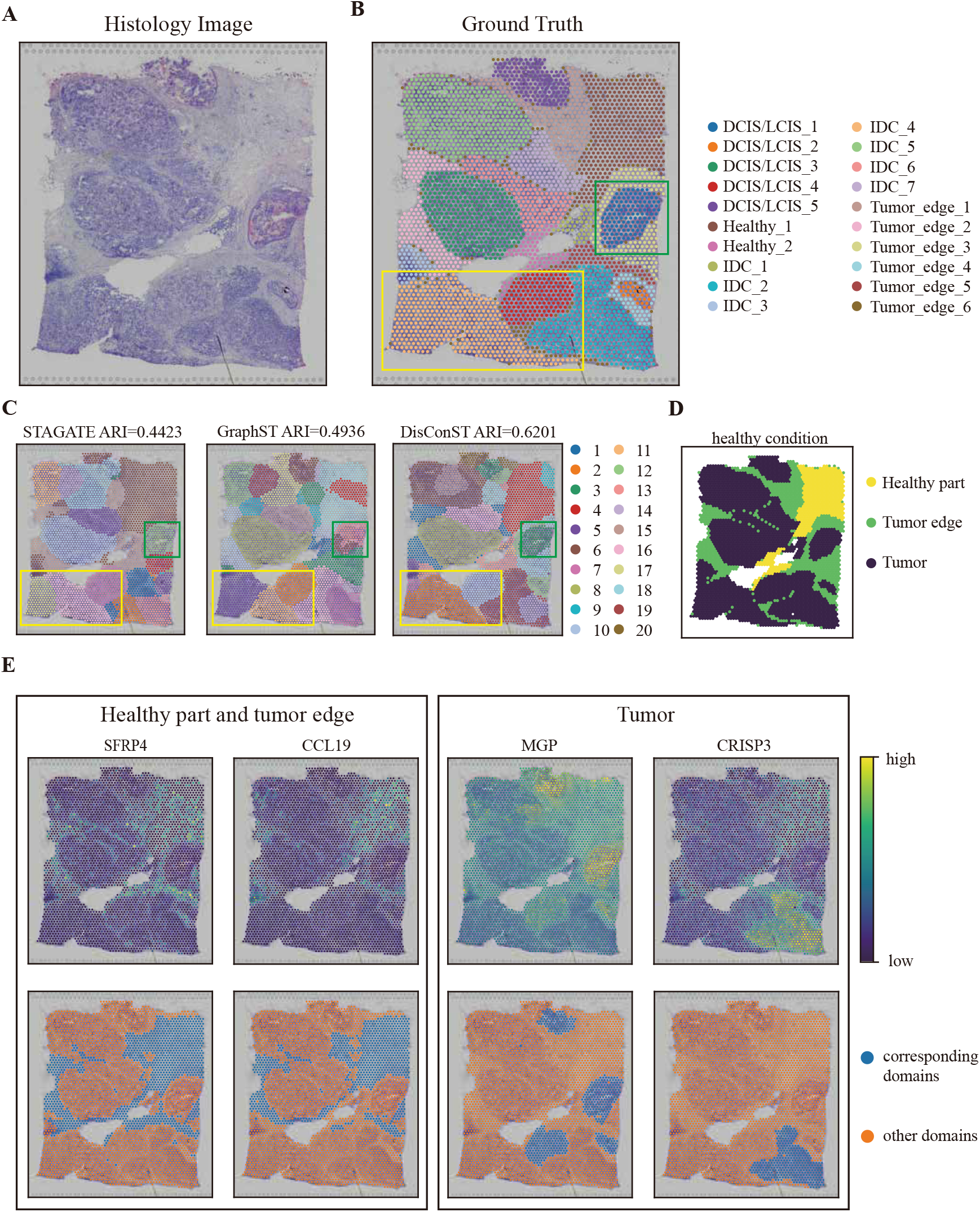
DisConST effectively dissects the immune microenvironment. **A**. Histological image of a human breast cancer slice. **B**. Ground truth annotations of the breast cancer slice based on H&E staining. **C**. Spatial domains identified by STAGATE, GraphST, and DisConST in the breast cancer tissue. **D**. Healthy regions of the breast cancer slice, annotated according to the ground truth. **E**. Genes highly

To further demonstrate our ability to distinguish between diseased and healthy areas, we validated our spatial domain identification results using the spatial expression patterns of marker genes. As shown in Figure 6D, this slice was divided into three parts based on manual annotation, including the tumor area, tumor edge, and healthy area. Corresponding to these areas, we selected four marker genes—*SFRP4* [31], *CCL19* [32], *MGP* [33], and *CRISP3* [34]—which are crucial in the development or resistance of breast cancer. *SFRP4* and *CCL19* are notably expressed at high levels in the healthy area and tumor edge, while *MGP* and *CRISP3* demonstrate heightened expression in the tumor region. As illustrated in Figure 6E, the expression patterns of *SFRP4* and *CCL19* align well with DisConST’s identification of domains corresponding to the healthy and tumor edge regions. The expression pattern of *MGP* correlates with domains 3, 10, and 12, which correspond to DCIS/LCIS in the ground truth. Similarly, *CRISP3* is associated with domains 14 and 19, corresponding to IDC in the ground truth. These findings highlight DisConST’s capability to accurately identify the distribution of both tumor and healthy regions. In conclusion, DisConST’s precise delineation of tumor structures and its effective differentiation between healthy and tumor areas confirm its robustness in analyzing the immune microenvironment.

## Discussion and Conclusion

Spatial transcriptomics (ST) has revolutionized the study of gene expression by offering unprecedented insights into the spatial organization of tissues. By providing spatially resolved gene expression data at spot or cellular levels, it enables researchers to explore tissue structure, biological development, and disease mechanisms. A critical task within ST is spatial domain identification—defining regions with similar gene expression patterns, which is essential for understanding tissue organization and functionality.

Despite the advances in ST, current methods for spatial domain identification often face challenges, particularly with integrating spatial and transcriptomic data, as well as handling high dropout rates in gene expression profiles. To address these limitations, we introduce DisConST (Distribution-aware Contrastive Learning for Spatial Transcriptomics), a novel deep learning method specifically designed to improve spatial domain detection. DisConST integrates spatial information, transcriptomic data, and cell-type proportions within spots, utilizing advanced optimization techniques including zero-inflated negative binomial (ZINB) distribution fitting and graph contrastive learning. These innovations allow DisConST to extract more informative latent representations, leading to superior accuracy in spatial domain recognition.

Through extensive validation on diverse datasets—ranging from tissues, organs, and embryos across various sequencing platforms, under both normal and diseased conditions—DisConST demonstrated consistently higher accuracy compared to existing state-of-the-art methods. Its robust performance highlights its ability to address key challenges in spatial transcriptomics research. Moreover, the practical utility of DisConST extends to a range of biological contexts, including advancing our understanding of tissue organization, embryonic development, and the tumor immune microenvironment.

In conclusion, DisConST represents a significant leap forward in the field of spatial transcriptomics, offering a powerful tool for accurate spatial domain identification through its innovative, distribution-aware contrastive learning approach. By addressing key limitations of existing methods, DisConST has the potential to enhance both fundamental biological research and translational applications in disease Studies.

## Authors’ contributions

T.W., X.W. and J.C. conceived the experiment(s), P.Z., X.W., and H.S. conducted the experiment(s), T.W., P.Z., X.W., J.H., Y.W., J.P., and X.S. analyzed the results. T.W., J.C. and P.Z. wrote and reviewed the manuscript.

## Acknowledgments

The authors thank the anonymous reviewers for their valuable suggestions.

