## Supplementary materials for "DisConST: Deciphering Spatial Domains Using Distribution-aware Contrastive Learning for Spatial Transcriptomics"

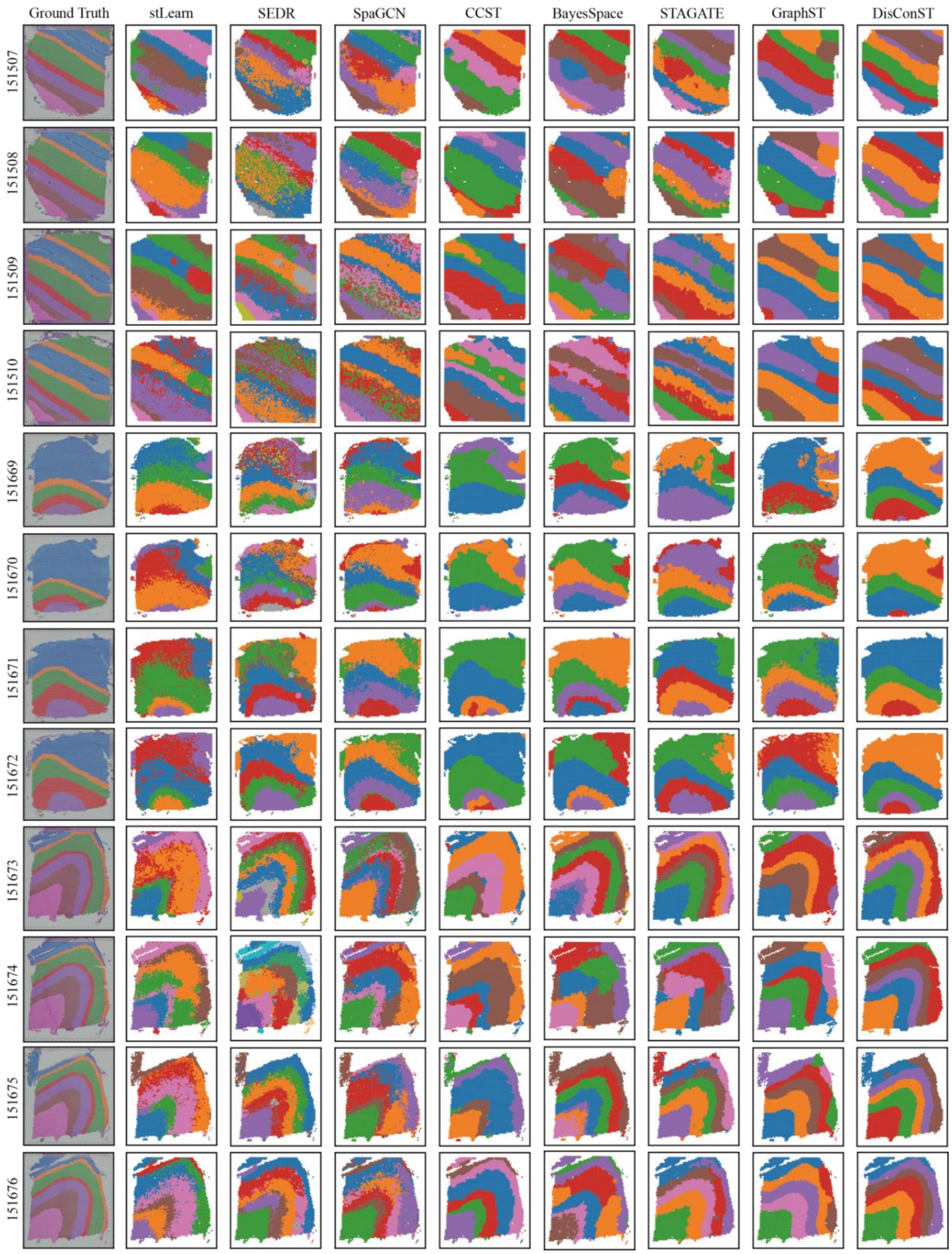


**Figure S1 Ground Truth and spatial domain identifications of eight methods on the 12 slices of DLPFC dataset, respectively**

*Note:* Those methods include stLearn, SEDR, SpaGCN, CCST, BayesSpace, STAGATE, GraphST, and DisConST.

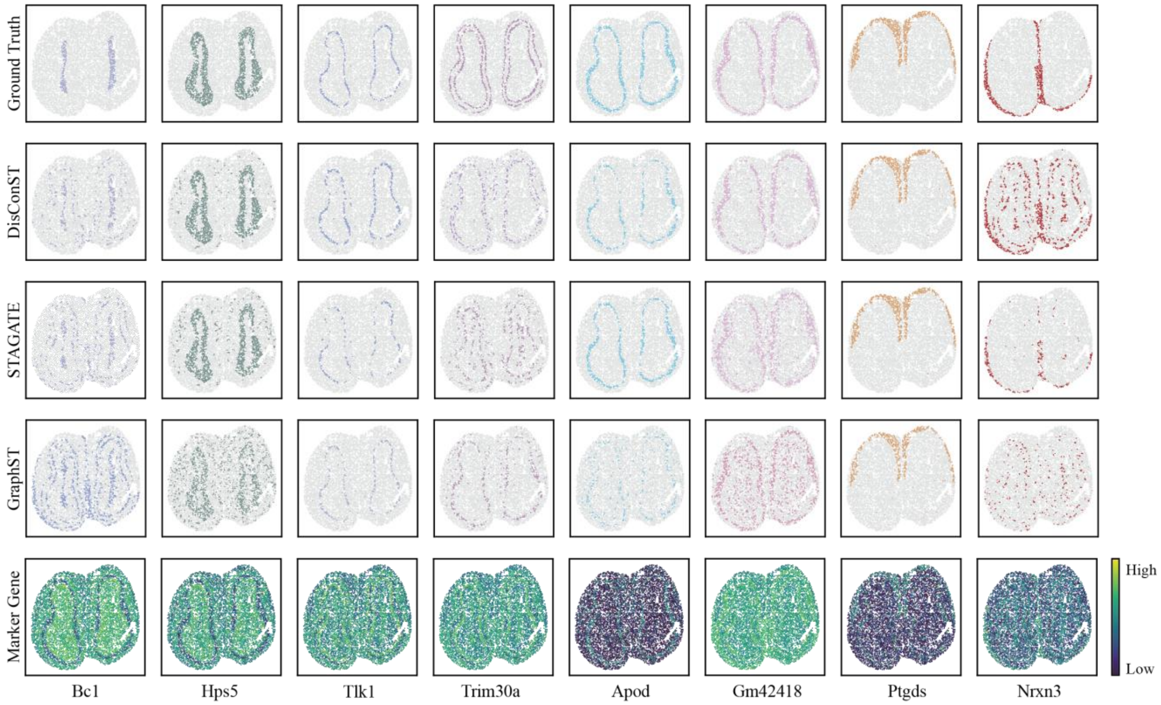


**Figure S2 Hierarchical structure of mouse olfactory bulb**

*Note*: The five line plots show the Ground Truth, identification results of DisConST, STAGATE, GraphST and corresponding marker genes of eight layers from the inner to the outer.

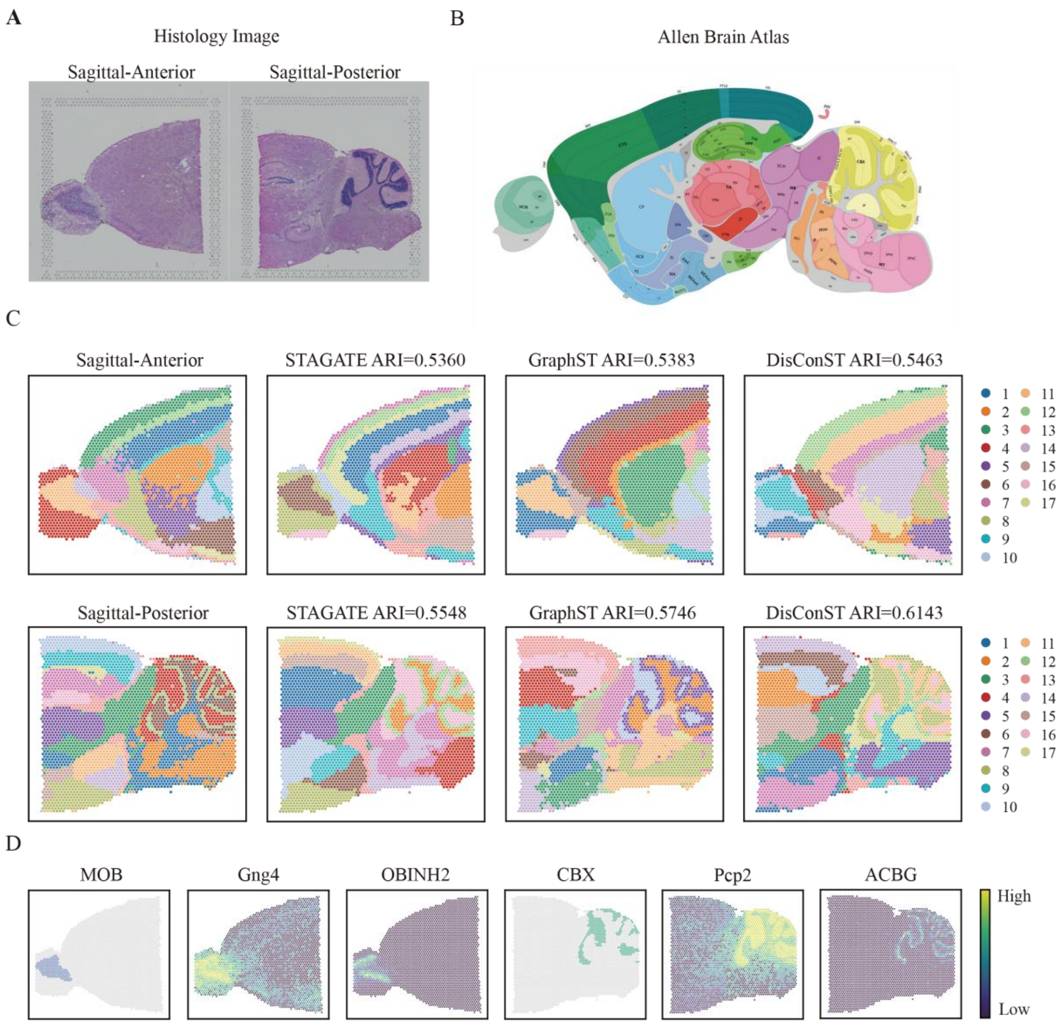


**Figure S3 DisConST accurately distinguishes different structures in the mouse brain**

**A.** The histology image of mouse brain sagittal-Anterior and sagittal-Posterior sections. **B.** The Allen brain atlas of the sagittal mouse brain in position 121. **C.** Visualization of spatial domain identification of three methods, STAGATE, GraphST, and DisConST in mouse brain sagittal-anterior and sagittal-posterior sections and corresponding Ground Truth. **D.** Marker genes and highly-distributed cell types in mouse olfactory bulb (MOB) and cerebellar cortex (CBX).

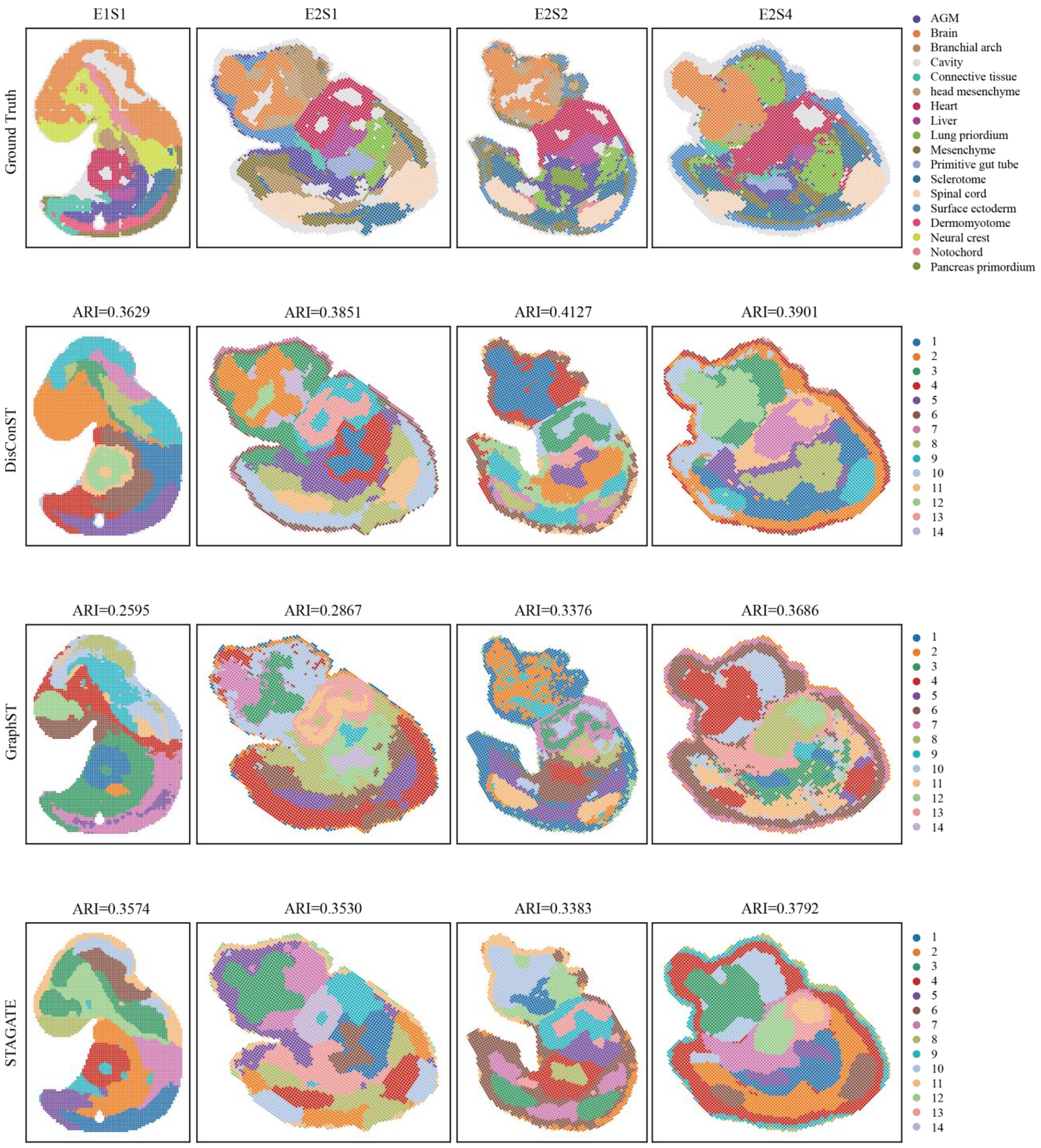


**Figure S4 Ground Truth and spatial domain identifications of STAGATE, GraphST, and DisConST on the Mouse Organogenesis Spatiotemporal Transcriptomic Atlas E9.5 data, respectively**

*Note*: E1S1, E2S1, E2S2 and E2S4 means different slices at stage E9.5.

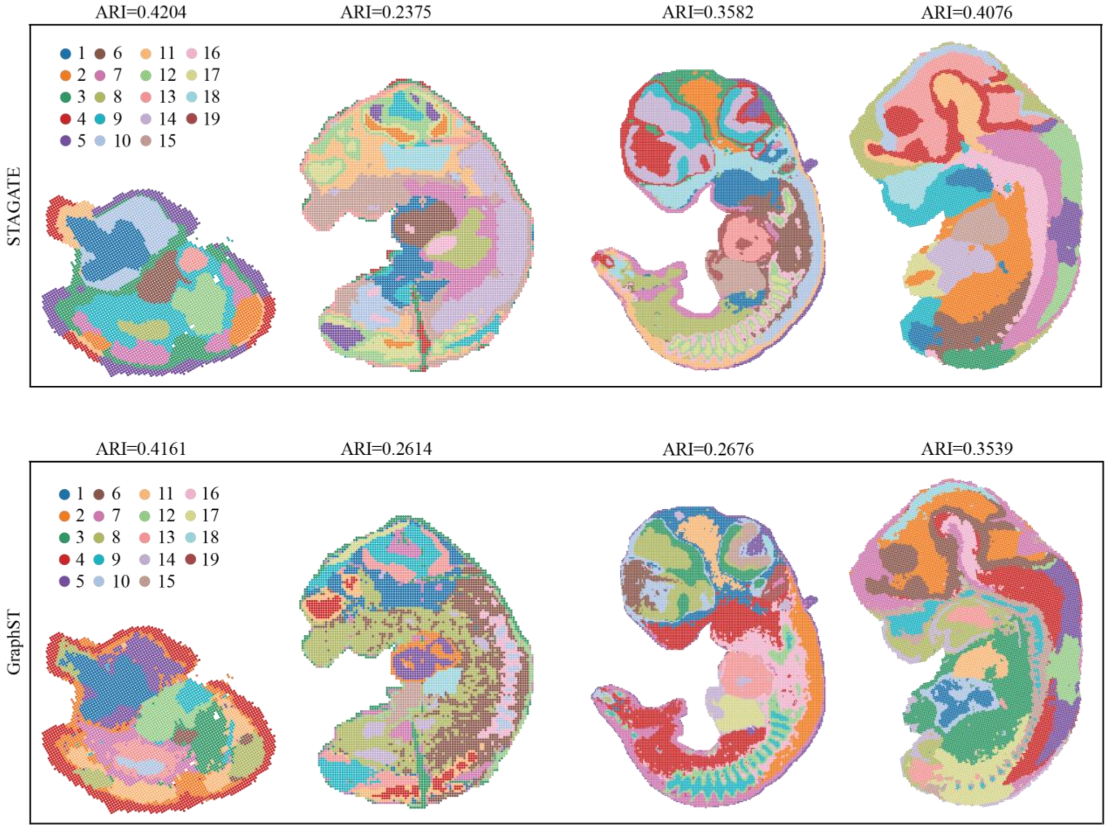


**Figure S5 Spatial domain identifications of STAGATE and GraphST on the 4 slices of mosta dataset (E9.5, E10.5, E11.5, E12.5), respectively**

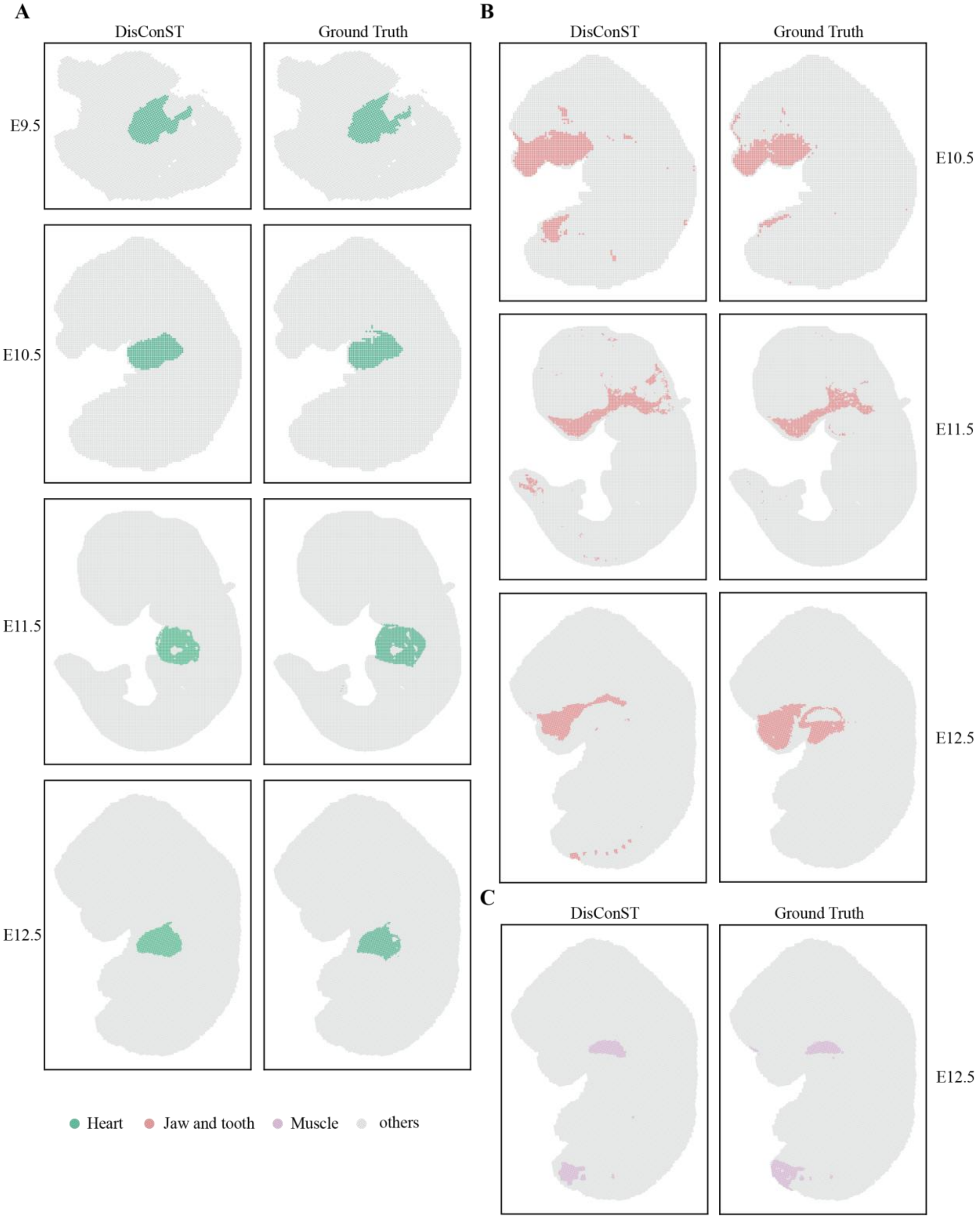


**Figure S6  The domain of heart, jaw and tooth and muscle**

**A.** Ground Truth and spatial domain identifications of heart among all four stages. **B.** Ground Truth and spatial domain identifications of jaw and tooth among E10.5, E11.5 and E12.5. **C.** Ground Truth and spatial domain identification of muscle on the E12.5 slice.

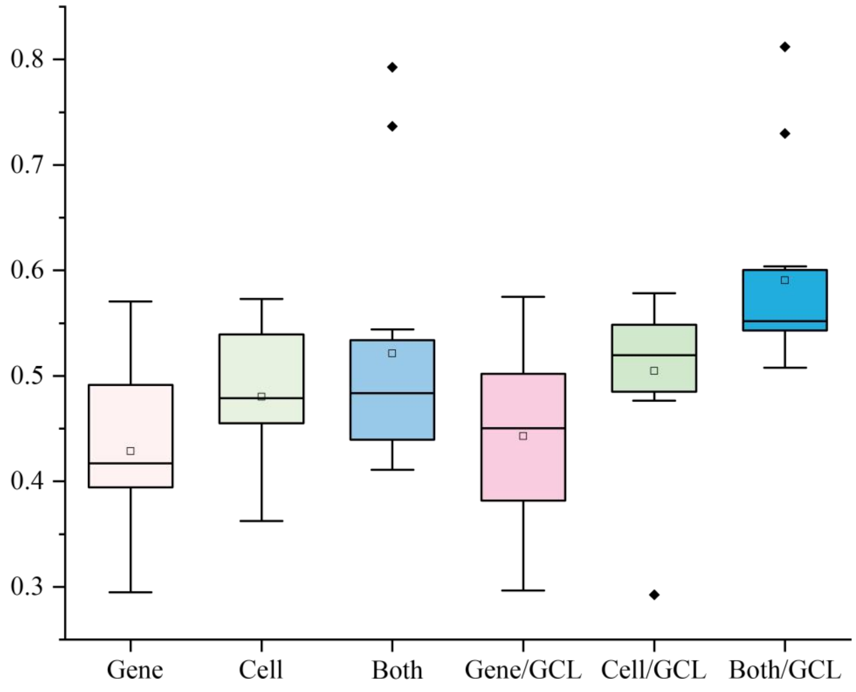


**Figure S7 The boxplot of DisConST ablation experiment on the 12 slices of DLPFC dataset**

*Note*: “Gene” means DisConST only uses gene expression data, “Cell” means using cell type proportion data only, “both” means using these two data without contrast learning loss. On the contrary, “Gene/GCL,” “Cell/GCL” and “Both/GCL” mean using these data with contrast learning loss. The boxplot’s small square, center line, box limits, and whiskers denote the average, median, upper and lower quartiles, and 1.5× interquartile range, respectively. These ARI scores are calculated using unrefined labels.

**Table 1 ARI scores of DisConST and seven comparison methods on 12 DLPFC slices**

| Slice | stLearn | SEDR | SpaGCN | CCST | BayesSpace | STAGATE | GraphST | DisConST |
|---|---|---|---|---|---|---|---|---|
| 151507 | 0.4556 | 0.4040 | 0.3798 | 0.4833 | 0.4690 | 0.5347 | 0.4575 | 0.5977 |
| 151508 | 0.2948 | 0.2560 | 0.4426 | 0.3097 | 0.4369 | 0.4919 | 0.4656 | 0.5531 |
| 151509 | 0.4148 | 0.2884 | 0.4566 | 0.4027 | 0.3814 | 0.4837 | 0.4746 | 0.5578 |
| 151510 | 0.2703 | 0.2867 | 0.4524 | 0.4332 | 0.3767 | 0.4800 | 0.4780 | 0.5763 |
| 151669 | 0.3399 | 0.2136 | 0.2920 | 0.2627 | 0.4704 | 0.2565 | 0.3983 | 0.5862 |
| 151670 | 0.1871 | 0.2033 | 0.3395 | 0.2810 | 0.4291 | 0.3902 | 0.4023 | 0.5611 |
| 151671 | 0.2810 | 0.4238 | 0.5268 | 0.6545 | 0.7334 | 0.5855 | 0.5835 | 0.8569 |
| 151672 | 0.3441 | 0.5013 | 0.5394 | 0.6217 | 0.4389 | 0.5994 | 0.5978 | 0.7639 |
| 151673 | 0.3133 | 0.4507 | 0.4513 | 0.5638 | 0.5499 | 0.6047 | 0.6164 | 0.6183 |
| 151674 | 0.3205 | 0.3613 | 0.4390 | 0.4003 | 0.2959 | 0.4407 | 0.5812 | 0.6542 |
| 151675 | 0.4502 | 0.4957 | 0.3152 | 0.3700 | 0.5297 | 0.5841 | 0.5362 | 0.5636 |
| 151676 | 0.3588 | 0.4654 | 0.3984 | 0.4016 | 0.3642 | 0.5518 | 0.5383 | 0.6070 |
| average | 0.3359 | 0.3625 | 0.4192 | 0.4320 | 0.4563 | 0.5003 | 0.5108 | 0.6247 |

**Table 2 The State-of-the-art methods for spatial domain identification in spatial transcriptomics**

| Method | Clustering method | Use histology image | Source code link | Reference |
|---|---|---|---|---|
| stLearn | Leidon[1] | Yes | https://github.com/BiomedicalMachineLearning/stLearn | Pham *et al. bioRxiv*, 2020[2] |
| SEDR | Leidon | No | https://github.com/JinmiaoChenLab/SEDR | Chen *et. al. bioRxiv*, 2021[4] |
| SpaGCN | DEC[4] | Yes | https://github.com/jianhuupenn/SpaGCN | Hu *et al. Nat. Methods*, 2021[5] |
| CCST | K-means[6] | No | https://github.com/xiaoyeye/CCST | Li *et al. Nat. Comput. Sci.*, 2022[7] |
| BayesSpace | mclust[8] | No | https://github.com/edward130603/BayesSpace | Zhao *er al. Nat. Biotechnol.*, 2021[9] |
| STAGATE | mclust | No | https://github.com/QIFEIDKN/STAGATE_pyG | Dong and Zhang, *Nat. Commun.*, 2022[10] |
| GraphST | mclust | No | https://github.com/JinmiaoChenLab/GraphST | Long *et al. Nat. Commun.*, 2023[11] |

**Table 3 Summary of all spatial transcriptomics datasets used for experiments in the work**

| Source | Section ID | #Spots | #Genes | #Clusters | Sequencing technology | Corresponding figures |
|---|---|---|---|---|---|---|
| Human dorsolateral prefrontal Cortex (DLPFC)[12] | 151507 | 4226 | 33538 | 7 | 10X Visium | Fig.2, Fig.S1, Fig.S6, Table S1 |
| | 151508 | 4384 | | 7 | | |
| | 151509 | 4789 | | 7 | | |
| | 151510 | 4634 | | 7 | | |
| | 151669 | 3661 | | 5 | | |
| | 151670 | 3498 | | 5 | | |
| | 151671 | 4110 | | 5 | | |
| | 151672 | 4015 | | 5 | | |
| | 151673 | 3639 | | 7 | | |
| | 151674 | 3673 | | 7 | | |
| | 151675 | 3592 | | 7 | | |
| | 151676 | 3460 | | 7 | | |
| Mouse olfactory bulb | - | 10000 | 26145 | 8 | Stereo-seq | Fig.3A |
| | - | 282 | 16034 | 4 | Spatial Transcriptomics | |
| Mouse brain serial | Section1 Anterior | 2696 | 31053 | 20 | 10X Visium | Fig.4 |
| | Section1 Posterior | 3353 | | 17 | | |
| | Section2 Anterior | 2825 | | 17 | | Fig.S2 |

| | | | | | | |
|---|---|---|---|---|---|---|
| | Section2 Posterior | 3293 | | 17 | | |
| Human breast cancer | - | 3798 | 36601 | 20 | 10X Visium | Fig.5 |
| Mouse Organogenesis [13] | E9.5 E1S1 | 5913 | 25568 | 12 | Stereo-seq | Fig.S5 |
| | E9.5 E2S1 | 5292 | 23756 | 14 | | Fig.S5 |
| | E9.5 E2S2 | 4356 | 24107 | 13 | | Fig.S5 |
| | E9.5 E2S3 | 5059 | 24238 | 13 | | Fig.6 Fig.S3 Fig.S4 Fig.S5 |
| | E9.5 E2S4 | 5797 | 23398 | 13 | | Fig.S5 |
| | E10.5 E2S1 | 8494 | 22385 | 18 | | Fig.6 Fig.S3 Fig.S4 |
| | E11.5 E1S3 | 27455 | 25741 | 19 | | Fig.6 Fig.S3 Fig.S4 |
| | E12.5 E2S1 | 23640 | 26688 | 18 | | Fig.6 Fig.S3 Fig.S4 |
| | - | 19416 | 351 | 22 | seqFISH | Fig.S3B |

*Note*: # means the number of.

[11] Long Y, Ang KS, Li M, Chong KLK, Sethi R, Zhong C, et al. Spatially informed

clustering, integration, and deconvolution of spatial transcriptomics with GraphST. Nat. Commun. 2023;14(1):1155.

[12] Maynard KR, Collado - Torres L, Weber LM, Uytingco C, Barry BK, Williams SR, et al. Transcriptome-scale spatial gene expression in the human dorsolateral prefrontal cortex. Nat. Neurosci. 2021;24(3):425-36.

[13] Chen A, Liao S, Cheng M, Ma K, Wu L, Lai Y, et al. Spatiotemporal transcriptomic atlas of mouse organogenesis using DNA nanoball-patterned arrays. Cell 2022;185(10):1777-92.
